# Comprehensive PAM prediction for CRISPR-Cas systems reveals evidence for spacer sharing, preferred strand targeting and conserved links with CRISPR repeats

**DOI:** 10.1101/2021.05.04.442622

**Authors:** Jochem NA Vink, Jan HL Baijens, Stan JJ Brouns

## Abstract

The adaptive CRISPR-Cas immune system stores sequences from past invaders as spacers in CRISPR arrays and thereby provides direct evidence that links invaders to hosts. Mapping CRISPR spacers has revealed many aspects of CRISPR biology, including target requirements such as the protospacer adjacent motif (PAM). However, studies have so far been limited by a low number of mapped spacers in the database. By using vast metagenomic sequence databases, we mapped one third (∼70,000) of more than 200,000 unique CRISPR spacers from a variety of microbes, and derived a catalog of more than one hundred unique PAM sequences associated with specific CRISPR subtypes. These PAMs were further used to correctly assign the orientation of CRISPR arrays, revealing conserved patterns between the last nucleotides of the CRISPR repeat and PAM. From the curated CRISPR arrays dataset we could also deduce CRISPR subtype specific preferences for targeting either template or coding strand of open reading frames. While some DNA-targeting systems (e.g. Type I-E and Type II systems) prefer the template strand and avoid mRNA, other DNA- and RNA-targeting systems (i.e. Type I-A, I-B and Type III systems) prefer the coding strand and mRNA. In addition, we found large scale evidence that both CRISPR adaptation machinery and CRISPR arrays are shared between different CRISPR-Cas systems. This could lead to simultaneous DNA- and RNA targeting of invaders, which may be effective at combating mobile genetic invaders.

## Introduction

CRISPR, an adaptive immune system provides heritable defence in the form of spacers, which are short nucleic acid sequences (28-36 bp) obtained from previous encounters with mobile genetic elements (MGE). These are stored in the bacterial or archaeal chromosome in CRISPR arrays (Jackson et al., 2017). CRISPR arrays contain spacers flanked on both sides by repeat sequences (∼30 bp) and are transcribed as a single RNA, and subsequently processed into multiple crRNAs. crRNAs can be loaded into effector complexes formed by Cas proteins, that subsequently scan the cell for nucleic acid targets. Base pairing between the spacer and target nucleic acids (protospacer) allows the specific binding of effector complexes to targets, which are then destroyed (Brouns et al., 2008; Marraffini, 2015). CRISPR-Cas systems are widespread in bacteria and archaea, with 42% of bacterial and 85% of archaeal genomes containing a CRISPR system (Makarova et al., 2020).

Both acquisition of new spacers (CRISPR adaptation) and target inactivation (CRISPR interference) are carried out by specialized sets of Cas proteins. *Cas* genes likely have originated from Casposons (Krupovic et al., 2014), a family of self-replicating transposons, and have since evolved many new genes and gene variants (Makarova et al., 2020). Based on the evolutionary classification of their *cas* genes, there are two classes of CRISPR-Cas systems. Class I systems contain crRNA-effector complexes made up of multiple subunits, while effector complexes of Class II systems are encoded by a single *cas* gene (Makarova et al., 2020). The two classes are further divided into six types, where each type is further divided into subtypes. The different types and subtypes do not occur homogeneously in nature, with Class II systems being nearly exclusive to bacteria (Makarova et al., 2020). More than 95% of CRISPR systems found in complete genomes are one of the first three types: Type I, II or III (Pourcel et al., 2020).

CRISPR systems can be studied on a mechanistic or on a functional level. Mechanistic features describe how CRISPR systems are able to fulfil their role. The mechanisms through which CRISPR systems operate are diverse. For example, some CRISPR systems defend the cell by targeting DNA (e.g. Type I, II, IV and V), whereas other CRISPR types target invader RNA (e.g. Type III and VI) (Makarova et al., 2020). Another important mechanistic feature is the presence of a protospacer adjacent motif (PAM), which DNA-targeting systems require to differentiate self from non-self (Gleditzsch et al., 2019; Hale et al., 2009; Mojica et al., 2009). Furthermore, the PAM is an important feature in the target search process of DNA-targeting systems within the cell (Vink et al., 2020; Xue et al., 2017). This motif sequence flanking the crRNA-pairing site, between one and five nucleotides long, not only differs between subtypes, but can also differ between *cas* gene orthologs within the same subtype, for example Cas9 variants (Gasiunas et al., 2020).

Functional features describe what purposes CRISPR systems fulfil within the cell. There is evidence for some CRISPR functioning beyond adaptive immunity (Westra et al., 2014), however even within the context of an adaptive immune system, CRISPR systems can serve different roles (e.g. as a first line of defence, or as an activator of other immune system pathways). This can be a reason why 23% of genomes with CRISPR systems contain more than one subtype (Bernheim et al., 2020), in spite of their costs (Nobrega et al., 2020; Vale et al., 2015). There are preferred combinations of certain subtypes, suggesting that there is an added benefit of having a specific combination of different subtypes present in the cell. The added benefit might consist of cooperativity between systems by formation of different lines of defence, avoidance of type-specific CRISPR inhibition by MGE or coupling of abortive infections mechanisms (Bernheim et al., 2020; Hoikkala et al., 2021; Pawluk et al., 2017; Silas et al., 2017). On the other hand, some CRISPR systems are specialized to protect from certain invaders, which may require multiple co-occurring systems to be present in a single genome to protect from different types of invaders. Type IV systems that co-occur together with Type I systems primarily target plasmids (Pinilla-Redondo et al., 2020) and Type III systems have been shown to be able to target a class of phages that other Type I and V systems cannot (Malone et al., 2020; S. D. Mendoza et al., 2020), indicating that specialization in targets is a potential reason for co-occurrence of different subtypes. Through cooperation and specialization, co-occurring subtypes can function complementary.

The functional and mechanistic features described above have been demonstrated experimentally for several microbial model systems, and these are often of specific interest to applications such as genome-editing. High-throughput assays to identify the PAM of CRISPR systems have been developed, but remain laborious (Gasiunas et al., 2020; Walton et al., 2021). The full diversity of PAM and other mechanistic and functional features of CRISPR-Cas systems in nature remain understudied. To improve our knowledge on mechanistic and functional features of single and co-occurring CRISPR systems beyond the model organisms, we relied on vast metagenomic sequence databases to computationally find targets for spacers from diverse bacteria and archaea. We mapped a third of the unique spacers to a target in publicly available metagenome sequence databases. We used the flanking regions of found spacer targets to build an initial PAM catalog of more than a hundred unique PAMs, and for more than half of the spacers in CRISPRCasDB (Pourcel et al., 2020). This was then employed to assign the correct orientation of transcription of CRISPR arrays, giving access to target strand information of invaders, further improving PAM predictions, and uncovering conserved links between repeat ends and PAM. Through the quantification of the spacers targeting template or coding strands we found that the preference for one of these strands is subtype specific, and indicates that some DNA-targeting systems (Type I-E, Type II-A and Type II-C) avoid RNA while other DNA- and RNA-targeting systems preferentially target RNA (Type I-A, Type I-B and Type III systems). We found spacers in co-occurring CRISPR systems to be compatible with both PAM and strand requirements, indicating that they may be shared between systems and will lead to both DNA and RNA targeting. Lastly, we identified three categories of multi-effector compatible spacers, which meet the PAM and strand requirements of co-occurring DNA and RNA-targeting systems.

## Results

### Blast analysis finds matches for 32% of spacers from CRISPRCasDb

The first step in our analysis was to select a set of CRISPR spacers and find potential matches to these sequences in DNA sequence databases. To this end, we selected the previously described CRISPRCasDB, which contained all spacers from 4266 complete bacterial and archaeal genomes (Pourcel et al., 2020). The spacers from CRISPRCasDB were then mapped to sequences from the NCBI nucleotide database as well as metagenomic databases with high number of prokaryotic or their virus sequences. Matches between spacers and sequences from the databases were found using BLASTn (Altschul et al., 1990). The matches were then filtered using an optimized approach which increased the number of matches while keeping the false positives to a minimum (Methods, Supplementary figure 1A). As indication of the false positive rate, we determined that for the matches found in NCBI nucleotide database 1% were eukaryotic or eukaryotic virus sequences, with 10% of matches in prokaryotic viral sequences and the majority (88%) corresponding to prokaryotic genome sequences (Supplementary figure 1A). This specificity towards prokaryotic sequences in a database that contains predominantly (83%) eukaryotic sequences shows that even though false positive hits cannot be excluded, the false positive rate is low.

**Figure 1.**
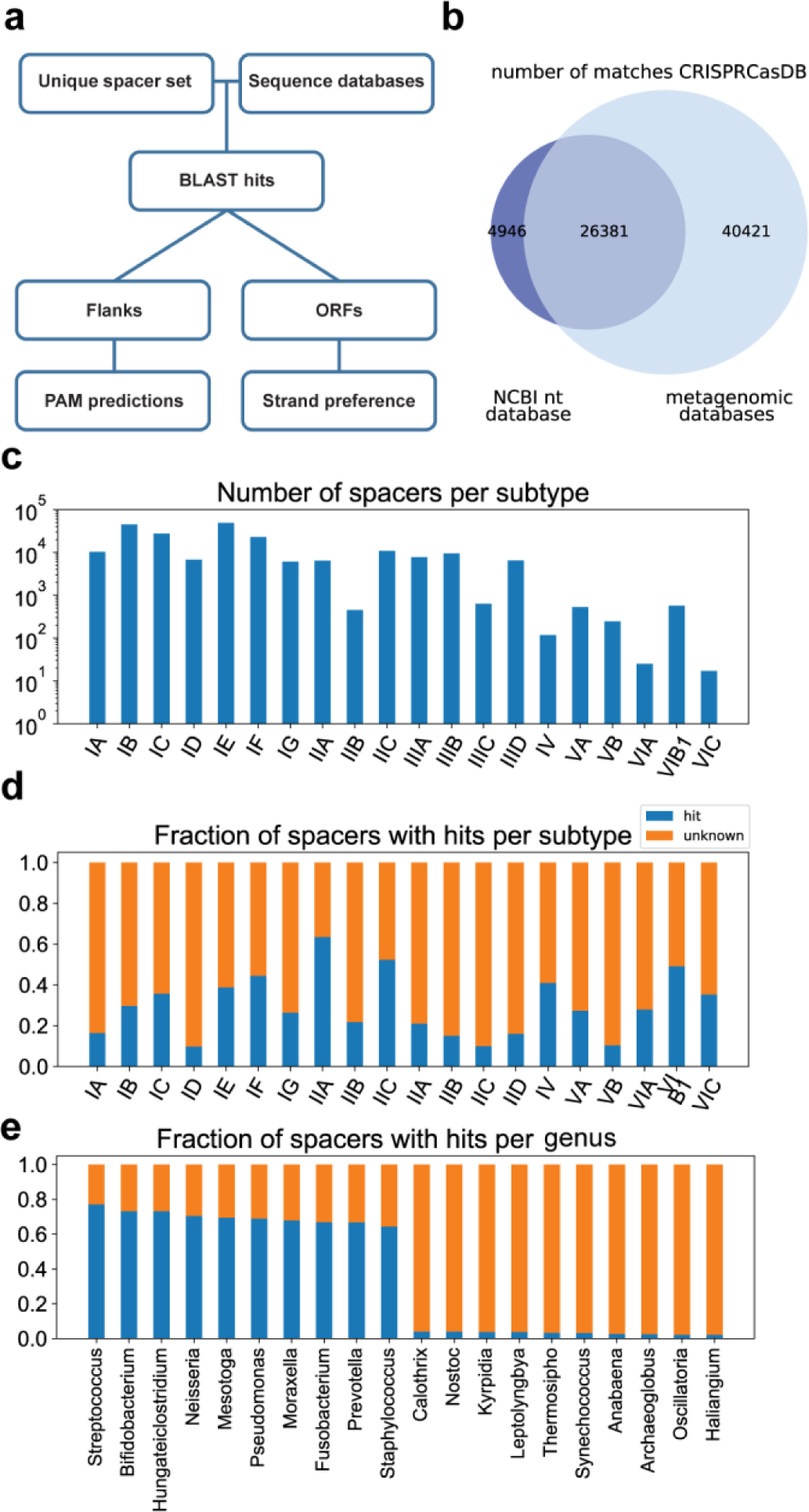
Spacer targets found with BLAST. (A) Computational pipeline for finding spacer targets. Targets of 72 099 spacers were found using blastn and filtered based on the fraction of spacer nucleotides matching a target sequence (See methods). (B) Venn diagram of spacers with matches in the NCBI nucleotide database versus metagenomic databases (C) Number of spacers per subtype. The subtype of a spacer was predicted based on similarity of the repeat sequence to repeats with a known subtype (See methods). (D) Fraction of spacers with hits per subtype. (E) Fraction of spacers with hits for the ten genera with the highest and ten genera with the lowest fraction of hits. Only genera with at least 500 spacers are shown.

From the 221,850 total unique spacers analysed, this optimized filtering approach resulted in 72,099 spacers (32% of total) with at least one match (Figure 1A), of which 31,327 spacers (15% of total) had a match in the NCBI nucleotide database (Figure 1B). The fraction of spacers with matches differed greatly between different genera, with *Streptococcus*, *Pseudomonas* and *Staphylococcus* among the genera with the highest fraction of matches (77%, 69% and 64% respectively) and *Calothrix*, *Nostoc* and *Thermosipho* among the lowest (4%, 4% and 3% respectively) (Figure 1C). Genera with high spacer matches typically occurred in well-sampled environments (human-associated), whereas the genera with lower matches occurred in what appear to be poorly sampled environments (soil, oceanic). A previous study (Shmakov et al., 2017) which looked for spacer matches in the NCBI nucleotide database found matches for 7% of spacers, using a more stringent 95% sequence identity and 95% coverage cut off as filtering thresholds. This difference in the fraction of spacers with matches in the NCBI nucleotide database indicates the added benefit and importance of our more sensitive filtering process. Additionally, the number of sequences in the database has increased in recent years from ∼230 billion to ∼700 billion bases. The most important factor for the increase in the number of spacers with matches however was the use of metagenomic databases, as the majority of unique spacer matches derived from these databases (Figure 1B, Supplementary Figure 1B).

To find the subtypes of the spacers, we aligned the CRISPR repeat sequences to repeat sequences with known subtypes, based on the method described by Bernheim et al., 2020. With the exception of subtype II-B for which we extracted 453 spacers, all analysed subtypes from Type I, II and III systems contained more than a thousand spacers (Figure 1D). An exceptionally high fraction of spacers with matches was found for subtypes II-A (63%) and II-C (53%), while subtype I-A, subtype I-D, and Type III subtypes had notably lower fractions of spacer matches than average (15%, 11% and 20% respectively). The differences in fractions of matches found between subtypes may be due to their phylogenetic distributions, where well-sampled genera have different subtypes than poorly sampled genera (see above). However, even within well-sampled genera the fraction of spacers with matches differs between subtypes, with Type III subtypes having fewer hits on average (22%) than other subtypes (38%). Overall, the large number of spacers with matches revealed sets of sequences that were targeted by each CRISPR-Cas subtype, which were then used to study mechanistic and functional aspects of CRISPR defence.

### Alignment of protospacer flanks reveals 114 unique subtype-specific PAMs covering 55% of spacers

One of the important mechanistic features of CRISPR defence for DNA targeting systems (type I, II, IV and VI) is PAM recognition (Deveau et al., 2008; Mojica et al., 2009; Shah et al., 2013). The first PAM was discovered in the alignment of bacteriophage sequences that were targeted by *Streptococcus* spacers (Bolotin et al., 2005). Later studies revealed more PAMs or the effect of mutant versions of the PAM (Anders et al., 2014; Fischer et al., 2012; Leenay et al., 2016; Musharova et al., 2019). We expand on these known PAMs that are limited to well-studied organisms by predicting new PAMs based on the alignment of the flanks of spacer matches (protospacers). The potential of this method for large scale PAM predictions was shown in a previous bioinformatics study (Mendoza & Trinh, 2018), with a key limiting factor being the number of spacers with matching targets. It was also previously shown that PAMs, acquisition machinery and repeat clusters co-evolve (Shah et al., 2013). We therefore increased the number of spacers with matches within one group by clustering spacers based on repeat similarity (>90% nucleotide identity and same repeat length). The sensitivity of PAM detection depends on the information content of the nucleotide positions of the PAM (signal) compared to the information content of the other flanking positions (noise). We found that clustering based on repeat similarity increased signal to noise ratio for PAM detection compared to clustering based on species-subtype (e.g. *Escherichia coli* I-E) or genus-subtype (e.g. *Pseudomonas* I-F). We furthermore found that spacers originating from organisms with very high or low GC-contents, displayed increased noise. We thus further increased the signal to noise ratio by adjusting the expected frequency of flanking nucleotides based on the average GC-content of the spacers within the cluster (Supplementary Figure 2). The flanks of unique hits within each cluster can subsequently be aligned, and with enough spacer hits, the information content reliably reveals the PAM sequence and position relative to the protospacer (Figure 2).

**Figure 2.**
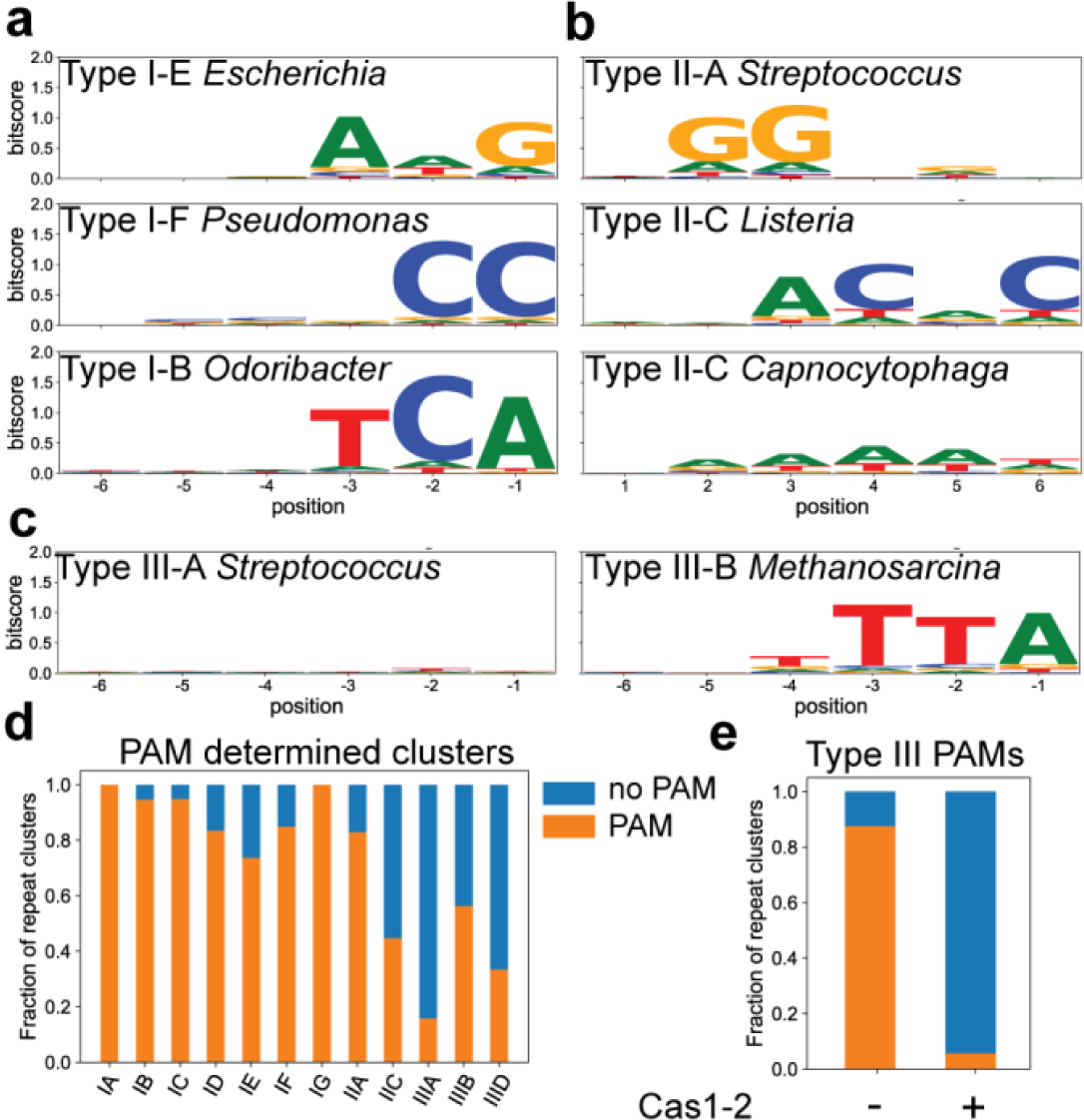
PAM determination of repeat clusters. (A) Sequence logos of upstream flank of hits to spacers from Type I repeat clusters. Sequence logos of protospacer flanking regions per repeat cluster. Y-axes show information content per nucleotide position. Label includes subtype of the repeat cluster and a representative genus in which this repeat cluster is found. (B) Same as (A) but for downstream flanks of spacers from Type II repeat clusters. (C) Same as (A) but for upstream flanks from Type III repeat clusters. (D) Frequency of PAM determined repeat clusters with more than 25 hits. Nucleotide positions were considered part of PAM with a bitscore of at least 0.4 and 10 times above the median bitscore of the 23 nucleotides surrounding the hits. PAM size was at least 2 nucleotides. (E) Frequency of PAM determined repeat clusters for Type III systems that contain Cas1-2 vs Type III systems that lack Cas1-2.

This clustering approach together with our large number of hits led to a PAM prediction for 123,144 spacers (55% of all spacers; Supplementary File 1 and 2). For Type I and Type IV the PAM is known to occur in the 5’ (upstream) flank of the protospacer, while Type II systems have their PAM in the 3’ (downstream) flank of the protospacer (Jackson et al., 2017) (Figure 2A). This well characterized feature of the PAM therefore allows the unique possibility to correctly orient CRISPR arrays given the rules described above. To measure the accuracy of CRISPR array orientation predictions, we compared predictions to experimentally determined orientations from a recent study using transcriptome sequencing (TOP) to determine the direction of transcription of arrays (Houenoussi et al., 2020). The 7968 experimentally inferred spacer orientations were the same as our predictions in 85% of cases, while only 33% of TOP predicted spacer orientations were the same as the CRISPRCasDb prediction (Supplementary data). We furthermore found that many Type I and Type III repeats for which we predicted the orientation based on the PAM, contained the 3’-end motif ATTGAAAC of their repeat (Supplementary Figure 3) described previously (Lange et al., 2013). This conserved motif is transcribed and forms the 5’ handle of the crRNA and is held by crRNA-effector complexes. Altogether, these findings indicate that the position of the PAM is a reliable indicator for the orientation of the CRISPR array, and can be used to annotate CRISPR array information, giving access to features such as spacer acquisition chronology and strandedness.

### Type I PAMs are shorter and closer to the protospacer than Type II PAMs

Sequence logos of alignments of Type I (Figure 2A) recover previously known PAMs including the subtype I-E AWG PAM found in *Escherichia* and subtype I-F CC PAM found in *Pseudomonas* (Leenay & Beisel, 2017), but also many previously undescribed PAMs. They are generally short (2-3 nt) and are well defined (high information content/bit score). For Type II PAMs, we found both short, well defined PAM motifs (such as *Streptococcus* II-A) as well as longer PAMs with less conserved PAM motifs (Figure 2B). Poorly conserved PAM motifs could be caused by a variation of PAMs used within the same repeat cluster or by the promiscuity of PAM recognition in Type II systems (Crawley et al., 2018). Additionally, many Type II PAMs consisted of multiple consecutive nucleotides of the same kind in a row, such as NAAAA (*Capnocytophaga* II-C). A low nucleotide conservation and repetitive nucleotide identity of a sequence motif can be caused by ambiguity in PAM distance to the protospacer, as this ambiguity will spread the nucleotide conservation over a larger range of positions from the protospacer. PAMs were found with the closest conserved nucleotide ranging 2 to 5 nucleotides away from the protospacer. The first nucleotide position on the 3’ end of the protospacer was always found to be an N for Type II PAMs. For a minority of subtype II-A and subtype II-C repeat clusters, a distinct lack of PAM was found (Supplementary figure 4B). As some Cas9 proteins from subtypes II-A and II-C can target RNA independently from a PAM sequence (Strutt et al., 2018), this RNA targeting could contribute to natural PAM-less variants that may inspire engineered PAM-less variants (Walton et al., 2020). Alternatively, PAM usage might be highly variable in these specific repeat clusters and could therefore obscure distinct PAM motifs. Overall, we found 114 unique PAMs (PAM-subtype combinations; Supplementary Data File 1), of which 43 PAMs in Type I systems (Supplementary Table 1), with each subtype containing at least two different PAMs and subtype I-B containing 12 different PAMs.

### 43% of Type III repeat clusters contain a PAM

Like the PAM-less Type II variants, some Type III repeat clusters were devoid of a PAM. This is expected, as RNA-targeting systems do not require a PAM to find a target (Figure 2C, Supplementary Figure 4), and rely on the Protospacer Flanking Sequence (PFS) to avoid self targeting (Deng et al., 2013; Elmore et al., 2016). Interestingly, other repeat clusters contained PAMs that appeared to be the same as Type I PAMs, which raised the question, why these clusters contained a PAM. We compared the PAM detection frequency for clusters with at least 25 unique spacer hits (Figure 2D). For Type I subtypes and subtype II-A, the majority of repeat clusters have a defined PAM, whereas for Type II-C and Type III systems the number of PAM-containing repeat clusters was lower, with Type III-A having the lowest (16%) and III-B the highest (56%) fraction of PAM-containing repeat clusters in Type III systems. As it was previously shown that Type III systems often lack their own acquisition machinery (Makarova et al., 2015), we hypothesized that the PAM found in Type III repeat clusters originates from the spacer acquisition machinery that Type I systems share with Type III systems. We observed that the PAM frequency in Type III clusters that lack their own acquisition machinery is high (95%; Figure 2E), whereas the PAM frequency is low in Type III clusters that contain their own *cas1*-*cas2* genes (8%). This supports the hypothesis that the PAM in Type III arrays originates from Type I spacer acquisition modules functioning in *trans*.

### Conserved patterns between PAM and repeats

PAMs usually differ from the ends of CRISPR repeats, which allows for self-nonself discrimination (Leenay et al., 2016; Mojica et al., 2009; Westra et al., 2013). Type III and other RNA-targeting CRISPR systems do not require a PAM, but many do require mismatching between the repeat end and the protospacer flanking sequence (PFS) (Johnson et al., 2019; Marraffini & Sontheimer, 2010). Given these previous observations, we wanted to investigate if there are conserved links between repeat ends and PAM of individual systems (Figure 3A), and whether Type III PAMs that originate from Type I spacer acquisition modules are also compatible with Type III PFS requirements.

**Figure 3.**
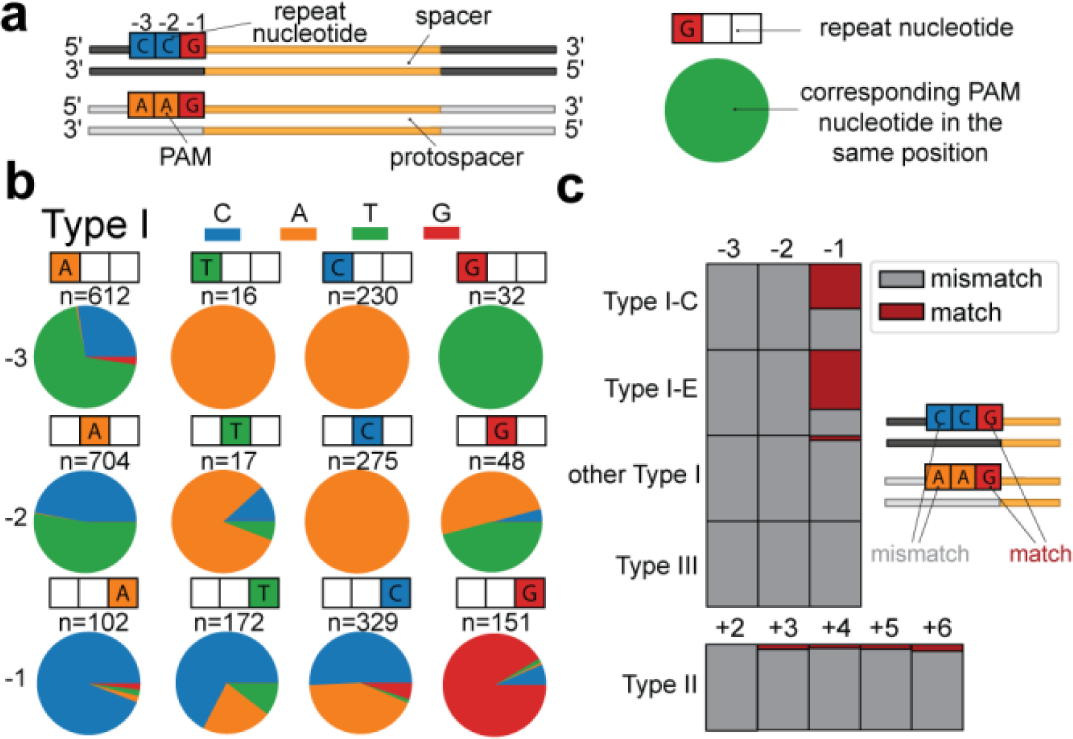
Relationship between repeat and PAM sequence. (A) Schematic of the analysis of PAM and repeat sequence. The nucleotide identity of the PAM in each position is compared to the nucleotide of the repeat. (B). PAM nucleotide frequency for Type I repeats. For each given repeat nucleotide position (indicated with coloured boxes) the PAM nucleotide (pie chart) for each unique PAM-repeat combination of our database. Number of occurrence is indicated above the pie chart (n). (C). The frequency of matches (red) and mismatches (grey) between the PAM and the corresponding repeat nucleotide for each position in relationship to the spacer. For Type II, the positions are compared on the other side of the spacer.

We collected all unique repeat-PAM sequence combinations in our dataset and compared the repeat nucleotide with the corresponding PAM nucleotide in each position. For Type I systems (Figure 3B) we found that the −3 and −2 nucleotide of the repeat can be a strong predictor of the corresponding PAM nucleotide, where a −3C in the repeat would lead to a −3A in the PAM, −3G to −3T, −3T to −3A. At the middle position a −2C would lead to a −2A in the PAM. (Figure 3B). The most common −2 and −3 repeat nucleotide is an A, in which case the PAM nucleotide mostly is either a T or a C. For the −1 position, the nucleotide identity of the PAM sequence cannot be predicted directly from the repeat sequence.

For Type II systems, most nucleotide positions can accommodate two or three PAM nucleotides (Supplementary Figure 5A). In +2 and +3 positions, the most common repeat nucleotide (T), accommodates either an A or G PAM nucleotide, which is analogous to the most common nucleotide in Type I systems (−3 and −2 adenine), which tends to co-occur with a C or T PAM nucleotide. For Type III systems, the variation of repeat nucleotides is smaller, but generally similar combinations are found as in Type I systems (Supplementary Figure 5B). Overall, the most conserved repeat-PAM co-occurrence patterns are found in the −2 and −3 positions of the Type I and Type III arrays.

These co-occurrence patterns suggest that in most cases the PAM that is used and selected for differs from the repeat. However previous studies have shown that in some cases, part of the repeat sequence is PAM-derived (Swarts et al., 2012). We then asked in what CRISPR subtypes the PAM matches the corresponding repeat nucleotide for each of the spacer flanking positions. When we counted the occurrence of a matching PAM, we found that this only occurred frequently in the −1 position of Type I-C (35%) and Type I-E (48%; Figure 3C). We found that these matches are associated with repeats that have TTC PAMs in Type I-C and AAG PAMs in Type I-E, which could indicate that the C of Type I-C repeat sequences is PAM-derived, as was similarly demonstrated for the G of AAG PAMs in Type I-E (Swarts et al., 2012).

In other positions and CRISPR types, >98% of the repeat-PAM combinations did not match each other, which shows that the general patterns between repeats and PAMs, and perhaps mechanism of self-vs nonself discrimination is conserved in all subtypes. In Type III systems all cases demonstrate mismatches between PAM and repeat, which is a requirement of functional Type III spacers (Johnson et al., 2019; Marraffini & Sontheimer, 2010). This finding demonstrates that the PAMs of Type III array spacers acquired with Type I acquisition modules are compatible with PFS requirements of Type III systems.

### Strand bias for the template or coding strand is subtype specific

Our method has revealed a large number of newly identified PAMs and has shown that Type III systems which lack their own acquisition machinery and co-occur with Type I systems, almost always contain a PAM. The presence of a PAM in these systems could enable Type I systems to use the spacers stored in Type III arrays as they are compatible with the PAM requirements of Type I effector complexes. Furthermore Type III effector complexes could benefit from a PAM-selecting acquisition module, as it excludes spacers with repeat-PAM matches (Figure 3C).

Besides the PFS, another requirement for type III spacers is that the spacer comes from the correct strand, as these complexes can only bind to the RNA transcripts. We wondered whether some species indeed use Type I and III dual functionality CRISPR arrays, as PAM-dependent DNA targeting and PAM-independent mRNA targeting are not mutually exclusive. We therefore asked whether spacers of DNA-targeting systems are also compatible with Type III surveillance complexes, if they happened to be picked from the correct strand.

To determine the potential ability of crRNA to target RNA, we measured the strand bias by counting the spacers that targeted the coding or template strand of predicted open reading frames (ORFs) (Figure 4A). As spacers targeting the template strand are unable to base pair the transcribed RNA, the fraction of spacers targeting the coding strand serves as an estimate of the RNA targeting ability of the crRNA. For example, in *Moraxella* IIIB arrays, a significant bias for the coding strand was found (88%, p<e^-11^) (Figure 4B). This bias allows Type III effectors carrying crRNA from those spacers to bind to their target RNA. However, also I-C spacers in *Moraxella*, for whose effectors this is not strictly required, show significant bias for the coding strand (p<e^-3^), indicating a selection for RNA-targeting spacers.

**Figure 4.**
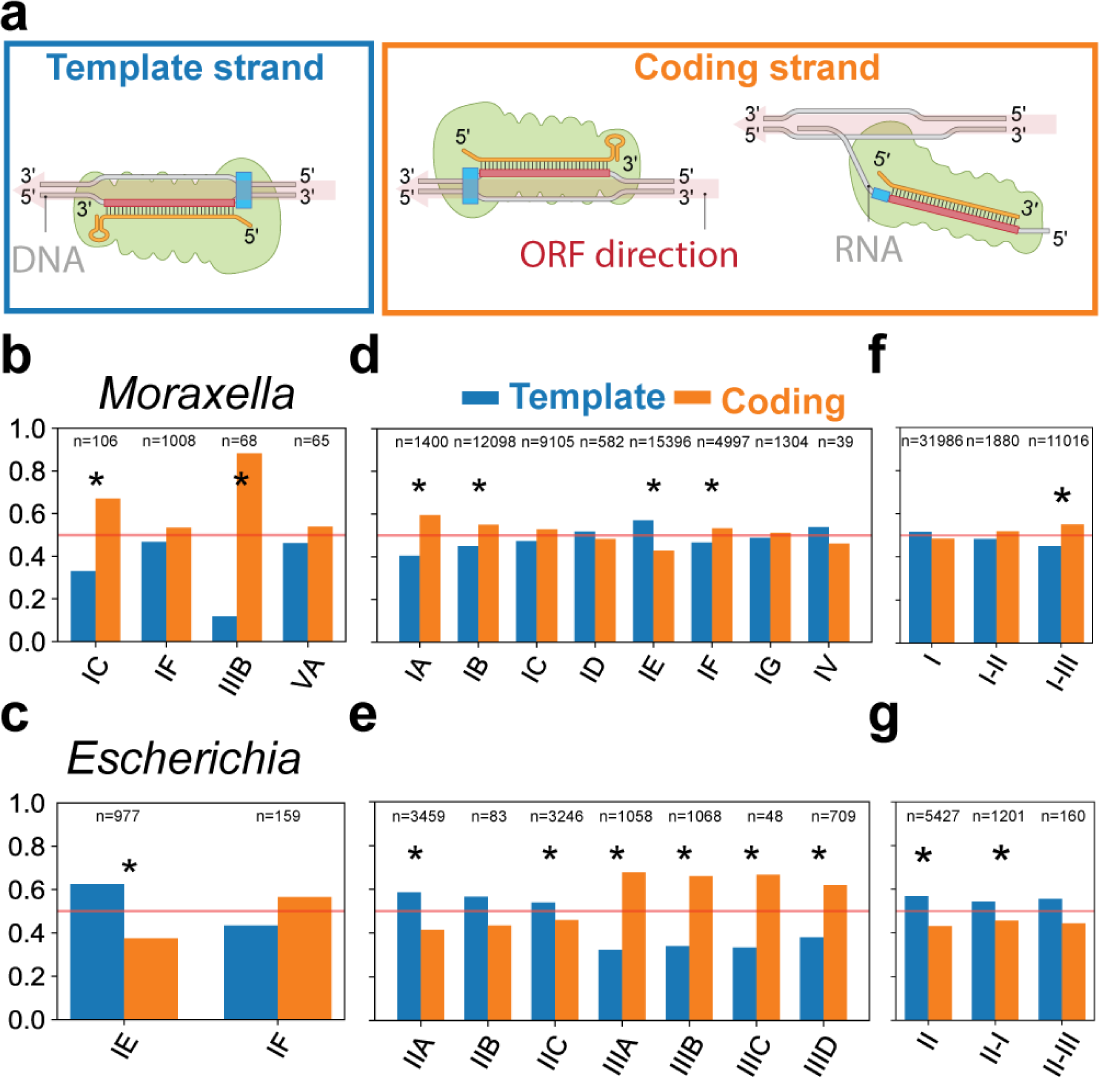
Template and coding strand targeting of spacers. (A) Schematic representation of a spacer targeting the template strand and a spacer targeting the coding strand inside an ORF. Spacers targeting the coding strand are also able to base pair with and target transcribed RNA. (B) Fraction of *Escherichia* spacers targeting template (blue) and coding (orange) strand by subtype. (C) Fraction of *Moraxella* spacers targeting template and coding strand by subtype. (D) Fraction of spacers targeting template and coding strand for Type I and Type IV subtypes. (E) Fraction of spacers targeting template and coding strand for Type II and Type III subtypes. (F) Fraction of spacers targeting template and coding strand for Type I. Spacers are grouped based on which other type of Cas effector genes are present in the genome. (G) Same as (F) but for Type II spacers. Significance of strand bias is calculated with a binomial test and a p-value<0.01 is indicated with an asterisk.

For *Escherichia* subtype I-E, 977 spacer matches inside ORFs were found, of which 611 (63%) targeted the template strand (Figure 4C), showing a significant bias for targeting the template strand (p<e^-14^) potentially avoiding RNA. No significant strand bias was found for *Escherichia* subtype I-F (43% template strand, p=0.11), suggesting that strand bias is CRISPR subtype specific.

Analysis of our complete dataset revealed general trends in the strand preferences for each subtype (Figure 4D, E). The strongest strand bias was found in Type III systems with an average of 65% of the spacers matching the coding strand (coding strand:template strand ∼ 2:1). This result demonstrates that there is selection in Type III systems for spacers to target the transcribed RNA. This selection can originate at the adaptation stage by dedicated adaptation machinery selecting from RNA/coding strands such as RT-Cas1 (Silas et al., 2016) or at the interference stage, where only functional RNA-targeting spacers are retained in the population (Artamonova et al., 2020). The strand biases we found are consistent with our curated CRISPR array orientation predictions, because an incorrect CRISPR array orientation prediction would obscure strand specific targeting. Type I-A and Type I-B also displayed significant strand bias for the coding strand although at lower levels (60% and 55%; p<e^-9^ and p<e^-14^ respectively).

Contrary to the Type III, Type I-A and I-B systems, we found a significant strand bias towards the template strand in in subtype I-E, Type IV and Type II systems, with the strongest bias found in subtype II-A (59%) and subtype I-E (57%). Given the high number of spacers in these groups the chance of observing this bias by chance is small (p <e^-23^ and p <e^-69^ respectively), again suggesting avoidance of RNA.

### Co-occurrence of Type I and Type III systems lead to PAM and strand targeting compatibility

As we noticed that Type III spacers were compatible with Type I PAMs in multiple cases, we next asked whether Type I spacers are compatible with RNA targeting in microbes with co-occurring Type I and III systems. We measured the strand bias of Type I spacers in genomes containing either combination of Type I, Type II and Type III surveillance complexes (Figure 4F). No significant strand bias was found for Type I spacers in the presence of Type I and/or Type II surveillance complexes. However, in the presence of Type I and Type III surveillance complexes, Type I spacers had a slight but significant coding strand bias (55%, p<e^-14^). This might be caused by increased selection pressure to keep RNA targeting spacers in the presence of RNA targeting surveillance complexes. This would suggest that spacers are selected to be compatible for both Type I and Type III effector complexes in such situations.

For Type II spacers, the presence of Type III did not significantly change the strand bias (Figure 4G). Given the natural tendency of Type II spacers to bias towards the template strand (Figure 4E), these findings suggest that Type II spacers are less compatible with co-occurring Type III effector complexes than Type I spacers.

### Three distinct categories of co-occurring multi-effector compatible arrays exist

The findings above indicate that subtype specific preferences exist for either the template or coding strand of the DNA. These preferences might enable or preclude compatibility between the spacers of co-occurring subtypes. We categorised all multi-effector compatible arrays that can be used by effector complexes from different subtypes. This means for co-occurring DNA-targeting systems these arrays need to have a PAM that can be used in both systems, whereas for co-occurrence of a DNA-target CRISPR-Cas system with an RNA targeting system, the arrays present in the genome need to both have the correct PAM and have a bias for the coding strand.

Overall, we can distinguish three main categories of co-occurring CRISPR-Cas systems in which spacers are compatible for multiple effectors (Figure 5A, Supplementary File 3).

**Figure 5.**
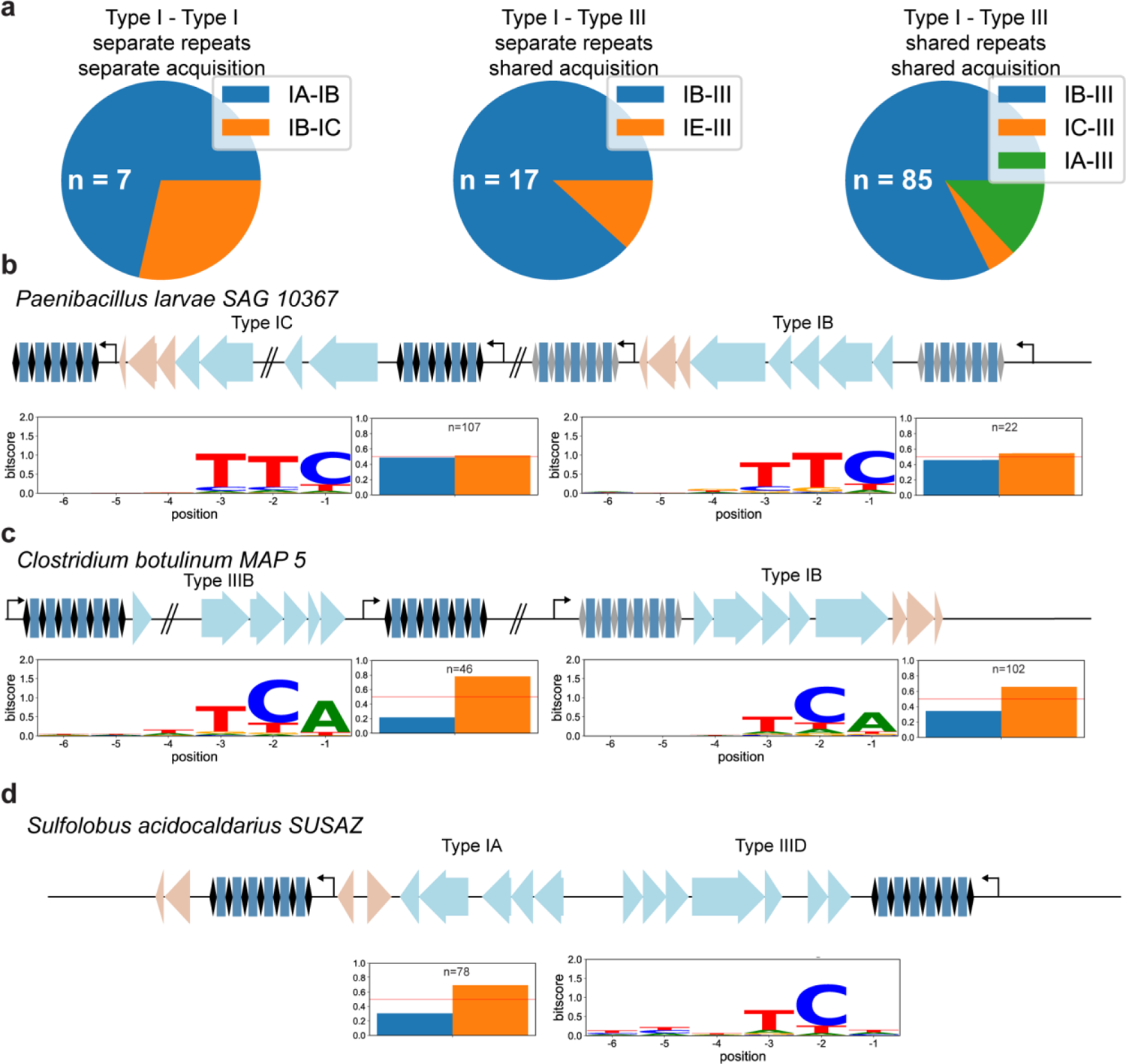
Different organisations of subtypes containing compatible spacer sequences. (A) Pie chart of frequency of genomes each category of organisation, based on the subtype combination involved. Total number of genomes for which this category was found (n) is noted in each chart (n).(B-D) Genome representations of examples for the different organisation categories, (b) Type I-Type I compatibility, (c) Type I-Type III compatibility (different repeat sequences), (d) Type I-Type III compatibility (same repeat sequences). Genes involved in interference (blue) and adaptation (red) are shown for the different subtypes within the genome. PAM logo and strand bias of each associated repeat cluster is depicted below the genomic representations.

The first category, exemplified by *Paenibacillus larvae* SAG 10367, consists of two co-occurring DNA-targeting systems which have their own adaptation machinery and their own repeat sequences. This is the smallest category and has been found in seven genomes (Figure 5A, B; Type I-A-Type I-B: 5, Type I-B-Type I-C: 2). We furthermore found 45 genomes of *Listeria*, which contain a Type I-B system and a Type II-A system from which the spacers have PAMs that might be compatible with both I-B and II-A effector complexes if the arrays are transcribed bi-directionally (CCN and NGG).

The second category, exemplified by *Clostridium botulinum* MAP 5, consists of a co-occurring DNA-targeting and RNA-targeting, with distinct repeat sequences but a commonly shared acquisition machinery (Figure 5A, C). We have only found evidence for multi-effector compatibility in co-occurring Type I and Type III systems. The Type I array in this category has a strand bias which indicates that the Type III effector complexes can use these spacers whereas the Type III arrays have the same PAM sequence as the Type I arrays allowing the Type I effector complexes to use these spacers. This category has been found in 17 genomes, mostly containing a Type I-B and a Type III system but also two genomes were found with a Type I-E and a Type III system.

The third category, exemplified by *Sulfolobus acidocaldarius* SUSAZ, consists of a co-occurring DNA-targeting and RNA-targeting system, with shared repeat sequences and shared acquisition machinery (Figure 5A, D). In this category the single repeat cluster present has a PAM and coding strand bias. This is the most common category of multi-effector compatible arrays which we detected in 85 genomes. It consists of co-occurring Type III systems with either a Type I-B (82%) but also Type I-A (13%) and Type I-C (5%).

Taken together, our data indicate that multi-effector compatible arrays are most prevalent between Type I and Type III systems. Within the Type I systems, the most common subtype to use multi-effector compatible arrays is Type I-B, but also Type I-A, Type I-C and Type I-E use these arrays. The Type III systems that use compatible arrays lack their own adaptation machinery, however repeat clusters in these co-occurring systems display a strand bias that suggests selection for RNA-targeting spacers. The information content is similarly strong for PAMs in Type III arrays as in Type I arrays, which demonstrates that the PAM is equally strong selected for Type I as shared Type III arrays.

## Discussion

In this study we have matched CRISPR spacers of complete genomes of bacteria and archaea with their targets in (meta)genome databases and subsequently analysed the genomic flanks of the protospacers. We computationally found targets for 32% of CRISPR spacers from thousands of bacterial and archaeal genomes. This is a major increase in spacer targets compared to previous studies and is due to our sensitive filtering process and use of metagenomic databases (Shmakov et al., 2017). We found that Type III spacers had the highest fraction of unknown targets of any CRISPR type. This was not solely caused by the phylogenetic or environmental occurrence of Type III systems, because the fraction of Type III spacers with unknown targets within a genus was typically higher than that of other types. This means that the targets of Type III systems are either under sampled, or that Type III spacers contain more mismatches to their targets, making them harder to find computationally. Recently, a single new study doubled the number of known RNA viruses including phages (Wolf et al., 2020), while another study greatly increased the number of known single-stranded RNA phages (Callanan et al., 2020), indicating that RNA phages have been poorly sampled. We predict the fraction of spacers with matches to increase with increasing numbers of available metagenomic data, especially including more RNA viruses and more data from poorly sampled environments.

By analysing the flanks of the spacer hits in great depth, we have generated a vast catalog of PAM sequences for each CRISPR repeat cluster. The repeat sequence is a good predictor of parts of the PAM sequence, and outperformed clustering based on genus-subtype classifications. This finding is corroborated by the position-wise comparison of PAM and repeat nucleotides, which shows certain repeat nucleotides predict PAM nucleotides. This may be helpful to either predict the PAM from scratch, or to further experimentally determine the PAM while reducing the degeneracy at certain positions, limiting the predicted PAM sequence space. The mismatch between repeat and PAM nucleotides generally holds, except for the Type I-E and Type I-C, where for some repeat clusters the repeat nucleotide matches the PAM at the −1 position. The most common PAMs of these systems (TTC for I-C; AAG for I-E) are also complementary to each other. These findings indicate Type I-C systems could have a similar mechanism of spacer acquisition with a PAM-derived last repeat nucleotide as in Type I-E (Swarts et al., 2012), even though these systems do not share related Cas1 proteins (Makarova et al., 2011) or repeat structures (Lange et al., 2013).

The PAM catalog can be used to predict the PAM for arrays in newly sequenced genomes and metagenomic contigs if they contain repeats that are closely related to the repeats in our database, which gives access to unexplored mechanistic and biotechnological potential. For repeats that are not in our database, the nucleotide identities of the repeat in the spacer flanking positions can be used to predict, with lesser certainty, which PAM it could have and select certain CRISPR systems of interest for further study.

Furthermore, the position of the PAM in the target is a reliable indicator for the orientation of transcription of CRISPR arrays. Correct prediction of transcription of CRISPR arrays gives access to measuring chronology of invader encounters and strand specific targeting of CRISPR-Cas systems, which is especially relevant for RNA targeting CRISPR. The spacers of Type III systems, which target RNA, have a bias towards targeting coding strands, making them capable of base pairing and thereby targeting RNA. Unexpectedly we also found several subtypes with a preference for the template strand (I-E and Type II). The reason for this type of strand bias is not yet clear, but we pose that this could be caused by a selection for spacers that do not target RNA (RNA avoidance), as DNA-targeting with these spacers might be impacted by inactivating complementary RNA (Jore et al., 2011). In addition, there might be a difference in binding or dislodging of crRNA effector complexes from the template strand vs coding strand by RNA polymerase (Clarke et al., 2018; Vink et al., 2020).

We have categorized multi-effector compatible CRISPR arrays whether they share the same repeats and/or acquisition machinery and whether only DNA, or both DNA and RNA are targeted. DNA-targeting systems that use multi-effector compatible arrays generally have their own acquisition machinery and the low frequency of this co-occurrence in nature might indicate that this is not actively selected for. It needs to be experimentally verified whether the spacers in these compatible arrays are actually shared between complexes. However, some crRNA sharing between DNA systems has already been observed experimentally, so it’s therefore likely to be found for more systems (Majumdar et al., 2015).

Multi-effector compatible arrays are much more common in co-occurring DNA- and RNA-targeting systems and the strand bias that occurs in Type I arrays indicates that Type III effector complexes are using these spacers and thereby creating selection pressure on the RNA binding potential of the transcribed crRNA. It also seems that the most commonly co-occurring Type I systems (I-A, I-B and I-C) that use compatible arrays, also have the largest coding strand bias. Whether this strand bias is induced by the presence of Type III or whether these subtypes by their nature have a strand preference and therefore became more commonly compatible with Type III systems is not yet clear. Interestingly, many of the subtype combinations that share PAMs also co-occur more often than expected by chance, suggesting they have positive epistatic interactions (Bernheim et al., 2020). Furthermore, repeat sequences of type I-A and I-B are in same repeat families with Type III repeats, providing further indications of their compatibility (Lange et al., 2013).

The experimentally determined spacer sharing in *Marinomonas mediterranea* (Silas et al., 2017) described previously does not fall within the categories in this study as the Type III system has its own adaptation machinery. In this case, the systems are not mutually compatible because the Type I systems cannot use the Type III spacers due to a lack of PAM, which we have not further investigated in this study. Also the other previously experimentally described spacer sharing systems in *Pyrococcus* (Majumdar et al., 2015) and *Flavobacterium* (Hoikkala et al., 2021) were not found due to a lack of sufficient hits, which demonstrates that these bio-informatic analysis likely underestimate the number of systems that can cooperate.

The discovery of multi-effector spacer compatibility in a large number of genomes in this study together with previous experimental evidence of spacer sharing of RNA and DNA-targeting systems (Deng et al., 2013; Majumdar et al., 2015; Silas et al., 2017) shows that there is selection pressure to share spacers cooperatively within arrays. The evolutionary benefits of such cooperativity could be profound. Firstly, as two subtypes generally have different mismatch tolerance (Anderson et al., 2015; Fineran et al., 2014; Manica et al., 2013), targeting the same sequence with two subtypes can reduce the probability of escape mutation. Secondly a combination of an RNA and DNA targeting systems can provide multiple layers of defence, where RNA-targeting might give more time for DNA-targeting systems to destroy the invader before the cell is taken over (Vink et al., 2020). Thirdly the length of arrays in a genome has recently been shown to be limited by auto-immunity (H. Chen et al., 2021). By sharing spacers, each subtype is supplied with a maximum diversity of spacers while self-targeting costs are minimized. Lastly the different mechanisms these systems use allows for complementary and distinct benefits. The priming mechanism (Datsenko et al., 2012; Nicholson et al., 2019), unique to DNA targeting systems can accelerate spacer acquisition for both systems, whereas cOA signaling pathways (Kazlauskiene et al., 2017; Niewoehner et al., 2017), unique to Type III, could activate defence systems that benefit both systems.

Altogether this study highlights the wealth of information that can be retrieved by analysing the targets of CRISPR spacers on a large scale. It furthermore demonstrates under what conditions CRISPR-Cas systems can cooperate and provides a large catalog of PAM predictions and targeted MGEs awaiting further study.

## Materials and Methods

### CRISPR spacers and sequence data

221 089 spacers along with information on *cas* gene presence, genome and repeat sequence were obtained from CRISPRCasDb (Pourcel et al., 2020) in February 2020 and the taxonomy of the genomes was obtained from NCBI Taxonomy database (Federhen, 2012). We created our own sequence database by combining all sequences from the NCBI nucleotide database (Benson et al., 2018; Pruitt et al., 2005), environmental nucleotide database (Sayers et al., 2009), PHASTER (Arndt et al., 2016), Mgnify (Mitchell et al., 2020), IMG/M (I. M. A. Chen et al., 2017), IMG/Vr (Paez-Espino et al., 2019), HuVirDb (Soto-Perez et al., 2019), HMP database (Peterson et al., 2009), and data from Pasolli et al., 2019. All databases were accessed in February 2020.

Subtypes were predicted based on the repeat sequences using the subtype predictions and method described by Bernheim et al., 2020, where the subtype of a spacer was inferred by the similarity of its repeat sequence to repeat sequences with known subtype (74% identity threshold to infer subtype).

### Blast hits and filtering

Hits between spacers and sequences from the aforementioned databases were obtained using the command line blastn program (Altschul et al., 1990) version 2.10.0, which was run with parameters word_size 10, gapopen 10, penalty 1 and an e-value cutoff of 1, to find as many potential targets as possible. These blast hits were then filtered to remove hits of spacers inside CRISPR arrays and false positive hits found by chance. Hits inside CRISPR arrays were detected by aligning the repeat sequence of the spacer to the flanking regions of the spacer hit (23 nucleotides on both sides). This alignment was done using the globalxs function from the Biopython pairwise2 package (Cock et al., 2009) with −3 gap open and −3 gap extend parameters. If more than 13 nucleotides were identical in the alignment of at least one flank, the hit was suspected to fall inside a CRISPR array and was filtered out.

To minimize the number of hits found by chance, we filtered hits based on the fraction of spacer nucleotides that hit the target sequence, as this metric considers both the sequence identity and the coverage of the spacer by the blast hit. In a first step, only hits with this fraction higher than 90% were kept. To find targets for even more spacers while keeping the number of false positives low, we included a second step where hits with a fraction higher than 80% were kept if another spacer from the same genus hit the same contig or genome in the first step. This second step did not introduce hits on any new contigs or genomes and was based on the assumption that multiple spacers from the same genus hitting the same contig or genome is unlikely to be caused by chance. Finally, we removed spacers that were shorter than 27 nucleotides (54 spacers) and removed 7 spacers that were hitting aspecifically, such as inside ribosomal RNAs or tRNAs. This left 72,099 unique spacers with target hits for downstream analysis.

### Protospacer flank alignment for orientation and PAM predictions

The PAM is known to occur on the 5’ end of the protospacer for Type I, Type IV and V CRISPR-Cas systems, and on the 3’ end for Type II systems (Collias & Beisel, 2021; Jackson et al., 2017). We used this property to predict the orientation of transcription of CRISPR arrays and sequence of crRNA. The PAM sides were compared to the nucleotide conservation in the flanking regions of the spacer hits and the spacer orientations were predicted such that the flank with the greater conservation matched the known PAM side.

To measure the nucleotide conservation in the flanking regions, data from multiple spacers was combined based on the subtype and repeat sequences of the spacers. Highly similar repeat sequences from the same subtype were clustered using CD-HIT (Fu et al., 2012) with a 90% identity threshold. We hypothesized that similar repeat sequences would be used in a similar orientation and would utilize the same PAM sequences, as coevolution of PAM, repeat and Cas1 and Cas2 sequences has been shown previously (Alkhnbashi et al., 2014; Lange et al., 2013). For each repeat cluster the flanking regions of the spacer hits were aligned. To equally weigh each spacer within the repeat cluster, irrespective of the number of blast hits, consensus flanks were obtained per spacer. These consensus flanks contained the most frequent nucleotide per position of the flanking regions. From the alignment of consensus flanks the nucleotide conservation, or information content, in each flank was calculated in bitscore (Schneider & Stephens, 1990) using the Sequence logo python package. We corrected for GC-content of the targeted sequences by calculating the expected occurrences of each nucleotide based on the GC-content of the spacer sequences. To minimize the number of orientation predictions based on little or noisy data, we only predicted the orientation for repeat clusters when the alignment of consensus flanks consisted of at least 10 unique protospacers. Furthermore, the information content of at least two positions was higher than 0.3 bitscore and higher than 5 times the median bitscore calculated from 23-nt flanks on both sides. These parameters were chosen as strictly as possible, while still yielding orientation predictions for the highest number of spacers.

Using the orientation predictions described above, we predicted the PAMs for each repeat cluster by checking which nucleotide positions were conserved. To minimize PAM predictions based on noise, we only predicted the PAM for repeat clusters where the alignment of consensus flanks consisted of at least 10 unique protospacers. A nucleotide position was predicted to be part of the PAM when higher than 0.5 bitscore and higher than 10 times the median bitscore. These parameters were chosen as strictly as possible, while maximizing the number of repeat clusters with PAM predictions and minimizing the number of unique PAMs predicted.

We subsequently categorized and counted multi-effector compatible spacers in the following ways. Firstly by an occurrence of multiple repeat clusters with different subtype classification that both contained the same PAM, for two DNA targeting clusters (category I) or a DNA and a RNA targeting cluster (category II). Secondly if multiple *cas* gene clusters from different subtypes were in the vicinity of a single repeat cluster and their genomes did not further contain other arrays linked to these *cas* gene clusters they were counted as a third category multi-effector compatible array.

### Coding versus template strand targeting analysis

For each spacer target inside an open reading frame (ORF), we determined if the spacer targets the coding (DNA and RNA) or template strand (DNA-only) during transcription. The ORFs and its orientation were predicted using Prodigal (Hyatt et al., 2010) for one target sequence per spacer. The target sequence of each spacer was selected as the longest hit sequence in the NCBI nucleotide database, excluding ‘other sequences’, or, if no such sequence was hit, the longest hit sequence in metagenomics database. Using our spacer orientation predictions for Type I, II and IV spacers, and the orientation predictions from CRISPRCasDb for the other spacers, we checked if the spacer target (blast hit orientation) was on the coding or template strand of the predicted ORF. To test for significant bias towards either the temperate or the coding strand, a two-sided tailed binomial test was performed with an expected probability of 0.5.

## Supporting information

Supplemmentary Files description

## Data and Material availability

The datasets on which the analysis is based have been submitted as Supplementary Files. Scripts to reproduce figures are available on request.

## Acknowledgements

The authors thank Christine Pourcel and Pierre-Albert Charbit for supplying the CRISPRCasDB in a spacer-based format and all members of the Brouns groups for input during group discussions. S.B. is supported by a Vici grant of the Netherlands Organisation for Scientific Research (VI.C.182.027; NWO).

## Author contributions

S.B. and J.V. conceived and supervised the project; J.V. gathered databases; J.V. and J.B. wrote analysis scripts; J.V., J.B. and S.B. wrote the manuscript.

## Supplementary figures

**Supplementary figure 1.**
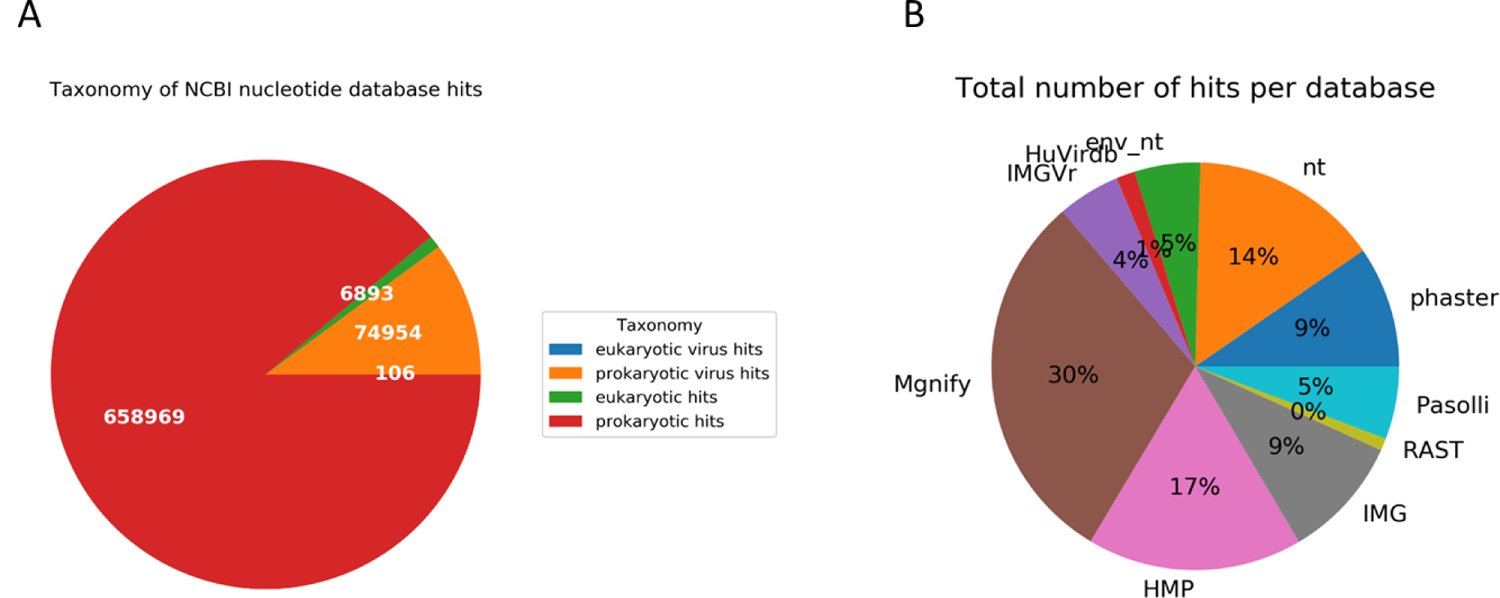
Taxonomy of spacer targets and number of found targets per database. (A) The taxonomy of targeted sequences of the NCBI nucleotide database was obtained from the NCBI taxonomy database. For hits in viral sequences, the taxonomy of known hosts was used to label the virus as a eukaryotic or prokaryotic virus. (B) The contribution of each database to the total number of hits after filtering. All databases were accessed in February 2020.

**Supplementary figure 2.**
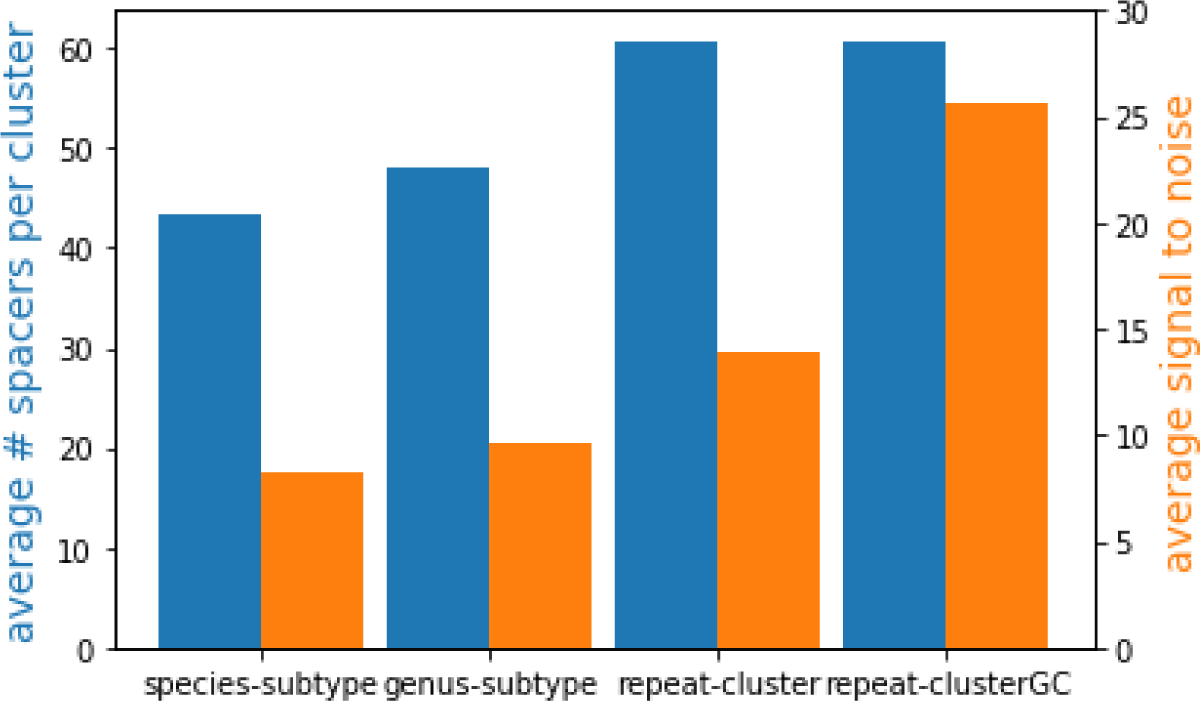
Average number and signal to noise ratio of clustered hits. Different clustering methods were compared for their average number of unique hits (blue) and average signal to noise ratio (orange). Signal to noise ratio was calculated by dividing the average information content of the two top positions in the flank (potential PAM nucleotides) by the median information content in sequence logos generated from flanks of hits. The clustering categories is based on whether spacers come from same species and subtype (species-subtype), from same genus and subtype (genus-subtype), from clusters of repeat sequences with 90% identity (repeat-cluster) or clusters of repeat sequences with 90% identity and additionally compensation for GC-content of spacers within the cluster (see Materials and Methods, repeat-clusterGC).

**Supplementary figure 3.**
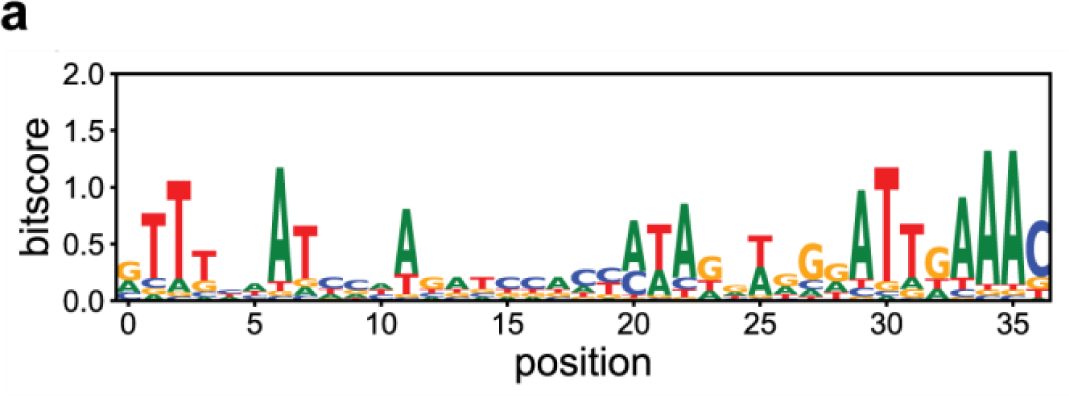
Sequence logo Type III repeats. ClustalW alignment of Type III repeats for which orientation was determined based on presence of PAM (n = 21 unique repeats). The 3’end of the repeat, which is the 5’ handle of the transcribed crRNA, has a conserved motif (ATTGAAAC).

**Supplementary figure 4.**
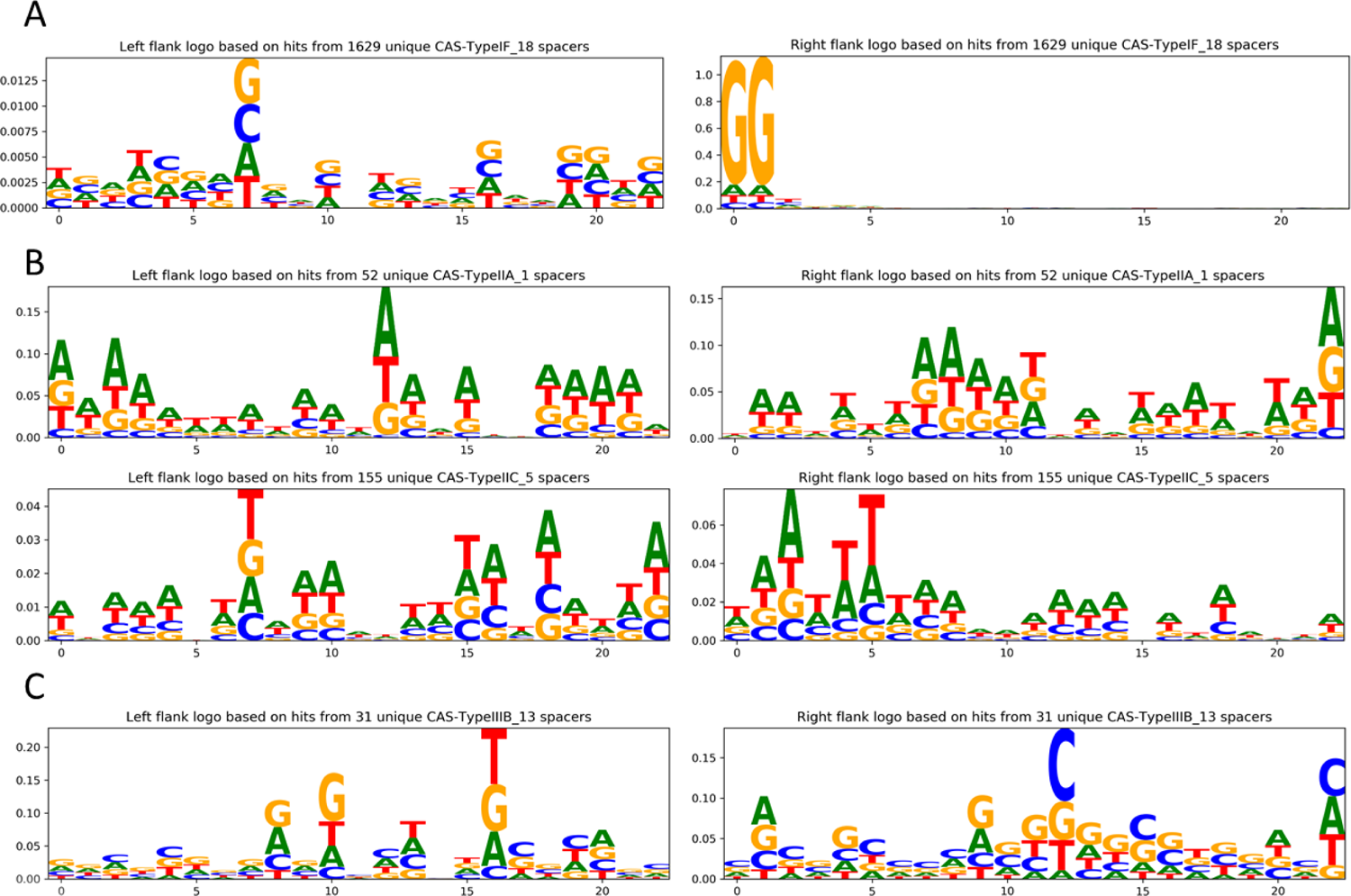
Sequence logos of protospacer flanking regions per repeat cluster. (A) The sequence logo of subtype IF repeat cluster is based on flanks of 1629 unique spacers. The reverse complement of the subtype IF CC PAM is found, due to incorrect orientation of spacers from the repeat cluster. (B) The sequence logos of subtype IIA and IIC repeat clusters. No positions with conserved nucleotides are visible, despite the high number of unique spacers for each cluster (52, 155 respectively). (C) The sequence logo of a subtype IIIB repeat cluster showing no positions with conserved nucleotides based on flanks of 31 unique spacers.

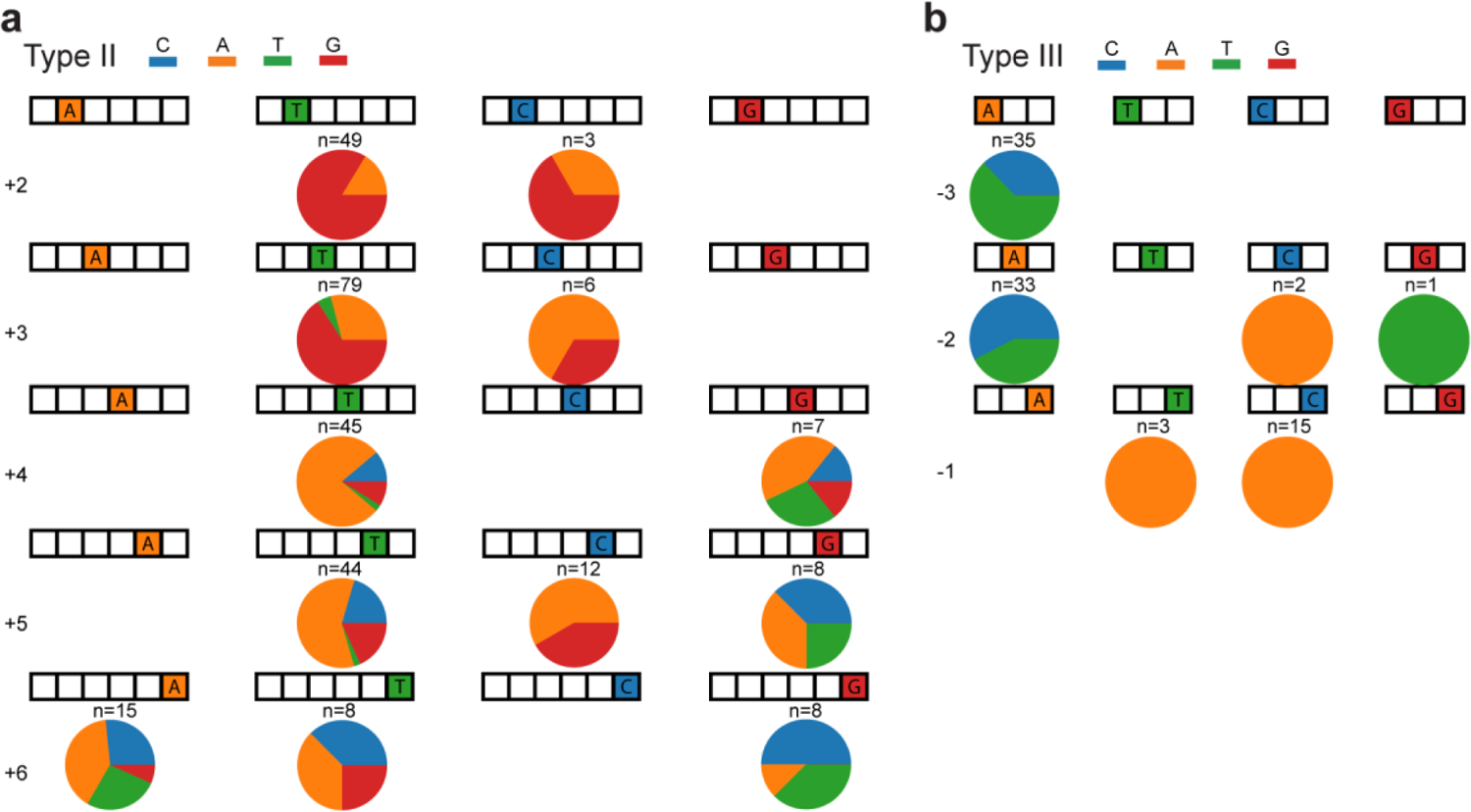
Relationship between repeat and PAM sequence of Type II and Type III systems. Same as Figure 3B except for Type II (A) and Type III (B) systems.

**Supplementary Table 1.**
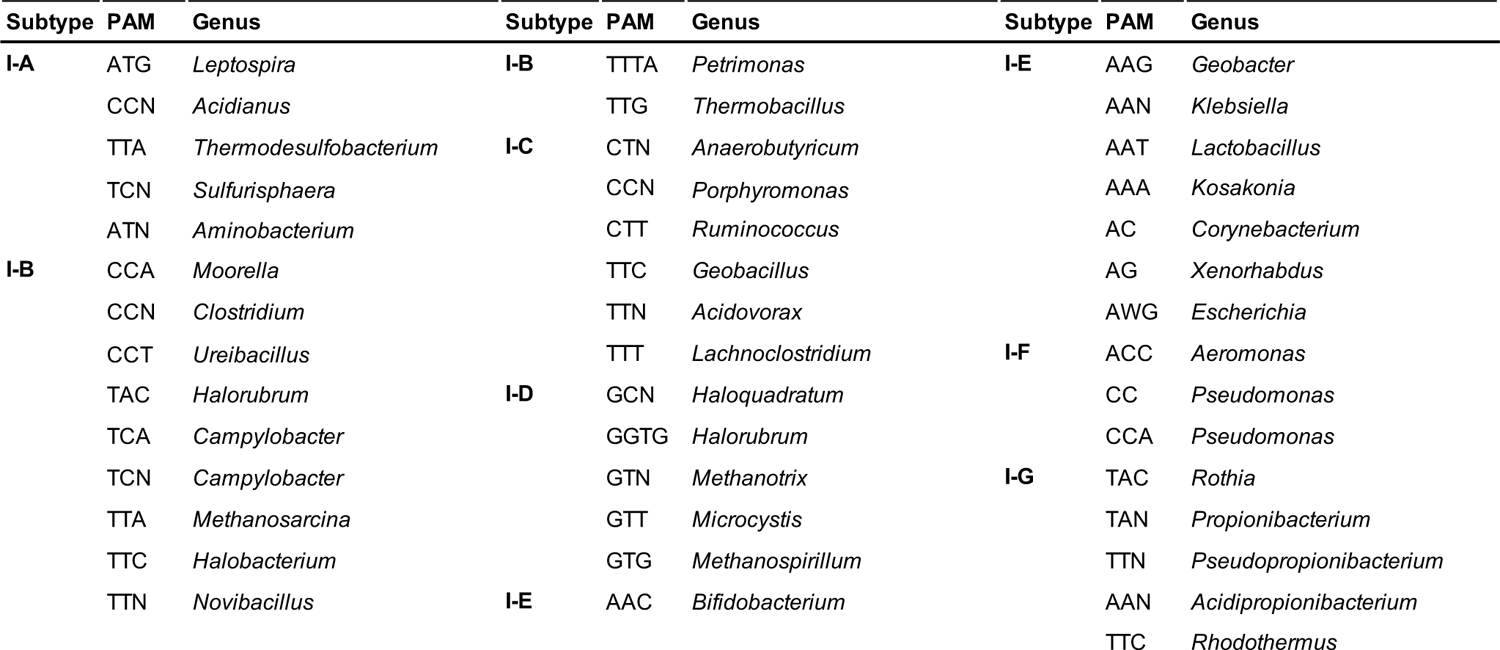
Unique Type I PAM sequences. Table of all unique Type I PAMs found for the different subtypes and representative genera that contain the repeat cluster for which each PAM was determined.

