## Supplementary material for "Comprehensive PAM prediction for CRISPR-Cas systems reveals evidence for spacer sharing, preferred strand targeting and conserved links with CRISPR repeats": Supplemmentary Files description

Supplementary files description

Supplementary file 1:

CSV file containing for each unique spacer in the CRISPRCasDB the following columns:

- **Spacers:** spacer sequence
- **Repeats:** repeat sequence in host(s) (can be multiple if multiple genomes contain same spacer)
- **Accessionnrs:** accession number of host(s)
- **Subtype:** subtype of array
- **cas_genes**: Cas_genes present in host(s)
- **hit**: if match found in (meta)genomic database equals 1 (else 0)
- **consensus**_flanks: consensus sequence of left and right flank from flanks of all the hits in databases to this spacer
- **repeat_cluster:** id of repeat cluster generated with CD-hit
- **strandbias:** Orientation of hit in reference to ORF (1 coding strand 0 template strand, -1 undetermined)
- **type:** Type of CRISPR array
- **orientation_CRISPRCasdb:** Orientation of spacer determined in CRISPRCasDB (Pourcel et al., 2020)
- **orientation_PAMbased:** Orientation of spacer determined in this study based on PAM
- **orientation_TOPbased:** Orientation of spacer determined with TOP (Houenoussi et al., 2020)
- **PAM:** PAM sequence of repeat cluster (if predicted)
- **Genus, Family, Order ….:** Taxonomy of host
- **Type I, TypeII, Type III…**: Whether host genomes contain genes related to specific Type (1 yes, 0 no)
- **Subtypesingenomes:** Which subtypes are in genomes
- **Subtypesinproximity:** Which subtypes are in proximity (<25000 bp from spacer)
- **Proximity_subtypes:** Distance of spacer to gene cluster of specific subtype
- **subtypesCas1**: Which subtypes are in genomes that contain a Cas1 protein

Supplementary file 2:

CSV containing PAM catalog (each unique repeat for which PAM was determined) with following columns: repeat, PAM and subtype

Supplementary file 3:

CSV containing genomes for which compatible arrays were found with following columns: accession number genome, compatible subtypes of array, PAM, category
